# Decoupled Surface Engineering Enables Cell-Interactive Edible Microcarriers across Food-Grade Polysaccharides

**DOI:** 10.64898/2026.11.07.765013

**Authors:** Minho Kim, Hwajin Oh, Chaeryeong Shim, Ahran Lim, Jeonghun Kim, Seung Soo Oh

## Abstract

Edible microcarriers for cultivated meat must support cell expansion while also contributing appropriately to the properties of the final food product. Food-grade polysaccharides exemplify this design challenge: despite their desirable food-related functions, they generally provide limited support for cell adhesion. This limitation calls for a strategy that introduces cell-interactive functionality without constraining bulk material selection. Here, we developed a decoupled surface-engineering strategy in which a poly-L-lysine/tannic acid (PLL/TA) interface is constructed on preformed polysaccharide microcarriers to enable cell-supportive functionality to emerge through interactions with the culture environment. On alginate microcarriers, PLL/TA modification markedly enhanced initial cell attachment, with 83% of modified microcarriers supporting attached cells compared with no detectable attachment on bare alginate. Cell-covered area subsequently reached 63.74% by Day 6, compared with 9.64% on PLL-coated microcarriers and negligible coverage on bare alginate. The PLL/TA interface promoted the recruitment of culture-medium proteins, including fibronectin, supporting an adaptive mechanism in which a cell-supportive interface develops through protein recruitment rather than predefined protein presentation. Importantly, the same surface modification enhanced early cell attachment when extended to pectin and carrageenan microcarriers and to Hanwoo bovine and Handon porcine muscle cells. On carrageenan microcarriers, the proportion of cell-bearing microcarriers increased from 30% to 100% for Hanwoo bovine cells and from 10% to 100% for Handon porcine cells following PLL/TA modification. These findings establish an adaptive and transferable surface-engineering strategy that enables cell-interactive functionality to be engineered separately from bulk material selection, expanding the design space for edible microcarriers in cultivated meat production.

## 1. Introduction

In cultivated meat production, edible microcarriers remain with the cultured cells as part of the final food product (Kawecki et al., 2024). Unlike conventional microcarriers that are removed after cell culture (Derakhti et al., 2019), they cannot be designed solely to provide an adherent environment for cell expansion. They must also satisfy food-related requirements, including edibility, texture, and nutritional value, because their material composition can directly influence the characteristics of the final product (Aguilera, 2019). In this regard, food-grade polysaccharides exemplify this challenge. Alginate, pectin, and carrageenan are widely used as gelling, thickening, and stabilizing agents in foods, making them attractive from a food-material perspective (Saha & Bhattacharya, 2010). However, these polysaccharides generally provide limited intrinsic support for the adhesion and growth of anchorage-dependent cells (Almeida et al., 2024; Lee & Mooney, 2012; Munarin et al., 2011). Their use as edible microcarriers therefore requires additional cell-interactive functionality without constraining the selection of the bulk material for its intended food-related role.

To improve the cell adhesiveness of edible microcarriers, previous studies have primarily relied on adhesion-supportive proteins. While some edible microcarriers have been fabricated using proteinaceous materials (Geurs et al., 2025; Norris et al., 2022), such proteins have also been incorporated during microcarrier fabrication (Jeong et al., 2026; Zernov et al., 2022) or directly coated onto preformed microcarriers before cell culture (Jaime-Rodríguez et al., 2025; Schlieker et al., 2026). Although these approaches have effectively supported cell attachment, incorporating proteins during fabrication changes the material composition and may consequently alter properties relevant to the final food product. Direct protein coating can avoid this problem by introducing cell-interactive functionality only at the surface. However, the resulting surface functionality is predetermined by the protein selected before cell culture, potentially requiring additional selection and optimization across different cell types and culture conditions (Bodiou et al., 2024). A more versatile strategy would therefore separate cell-interactive surface functionality from the underlying material composition without defining the biological interface in advance.

Here, we developed a decoupled surface-engineering strategy that allows food-relevant polysaccharide microcarriers to acquire cell-interactive functionality through adaptive interactions with the culture environment. We implemented this strategy by constructing a poly-L-lysine/tannic acid (PLL/TA) interface on preformed polysaccharide microcarriers, leveraging the multivalent noncovalent interactions of tannic acid with proteins (Jafari et al., 2022). PLL/TA modification transformed otherwise poorly adhesive polysaccharide surfaces into cell-supportive interfaces, with the proportion of cell-bearing microcarriers increasing from undetectable levels on bare alginate to 83% on PLL/TA-modified alginate in an MDA-MB-231 model system. The modified surfaces recruited proteins from the culture medium, indicating that a cell-supportive interface emerged adaptively during culture rather than through the predefined presentation of a specific adhesion-active protein. Importantly, the same PLL/TA modification promoted cell attachment across alginate, pectin, and carrageenan and was effective not only for the initial model cell system but also for Hanwoo (Korean native cattle) and Handon (Korean domestic pig) muscle cells, demonstrating its transferability across compositionally distinct food-grade polysaccharides and livestock-cell sources. Together, these findings establish an adaptive and transferable surface-engineering strategy that decouples cell-interactive functionality from bulk material composition, expanding the design space for edible microcarriers in cultivated meat production.

## 2. Materials and Methods

### 2.1. Materials

Alginic acid sodium salt (Sigma-Aldrich, St. Louis, MO, USA; Cat. No. 180947), low-methoxyl amidated pectin (SUN-IN Co., Ltd., Asan, Republic of Korea), and ι-carrageenan (TCI, Tokyo, Japan; Cat. No. C1805) were used for microcarrier fabrication. Calcium chloride dihydrate (Georgiachem, Suwanee, GA, USA; Cat. No. CC2091) was used for ionic crosslinking.

Poly-L-lysine (70,000-150,000 Da; Sigma-Aldrich, St. Louis, MO, USA; Cat. No. P6282) and tannic acid (Sigma-Aldrich, St. Louis, MO, USA; Cat. No. 403040) were used for layer-by-layer surface modification. 1M HEPES (T&I; Cat. No. BHE-9001), sodium chloride (Georgiachem, Suwanee, GA, USA; Cat. No. SC1010), Iron(Ⅲ) chloride (Sigma-Aldrich; Cat. No. 157740), and potassium hexacyanoferrate(Ⅲ) (SAMCHUN; Cat. No. P1825) were used for preparation of coating solutions and phenolic analysis.

Dulbecco’s Modified Eagle Medium (DMEM; WELGENE; Cat. No. LM 001-05), fetal bovine serum (FBS; EMD Millipore; Cat. No. TMS-013-BKR), penicillin–streptomycin (P/S; WELGENE; Cat. No. LS 202-01), and growth factor (SeaWith, South Korea) were used for cell culture. MDA-MB-231 cells were obtained from the Korean Cell Line Bank (KCLB; KCLB No. 30026). Hanwoo bovine muscle satellite cells (cell code: 790512) and Handon porcine muscle satellite cells (cell code: D24171) were provided by SeaWith Inc. (South Korea).

Calcein-AM (Thermo Fisher Scientific, Cat. No. C1430), propidium iodide (Sigma-Aldrich; Cat. No. P4864), and dimethyl sulfoxide (DMSO; Invitrogen; Cat. No. D12345) were used for live-cell fluorescence imaging. HiLyte™ 488-labeled fibronectin (FNR02; Cytoskeleton, Inc., Denver, CO, USA) was used for analysis of fibronectin association with microcarriers. Unless otherwise stated, deionized water was used for preparation of aqueous solutions.

### 2.2. Fabrication of polysaccharide microcarriers

Alginate, pectin, and carrageenan microcarriers were fabricated by spray-assisted ionic gelation. Sodium alginate was dissolved in deionized water at 2.5% (w/v) by heating on a hot plate set to 75 °C for approximately 20 min with intermittent manual mixing until completely dissolved. Pectin and carrageenan solutions were separately prepared at 1.0 % (w/v) in deionized water under the same conditions.

For microcarrier fabrication, 20 mL of polymer solution was loaded into a hand-operated spray bottle and sprayed into 200 mL of 20 mM CaCl₂ solution contained in a glass bath. The spray nozzle was positioned approximately 20 cm above the liquid surface and oriented vertically upward, away from the crosslinking bath. Spraying was performed at 5–10 s intervals, with gentle lateral shaking of the bath by hand between sprays. Spraying was continued until no further solution could be dispensed.

Fabrication time was recorded during spraying, and the solution flow rate was calculated by dividing the dispensed alginate solution volume by the fabrication time. Microcarrier yield was estimated by collecting an aliquot of the microcarrier suspension from the CaCl₂ crosslinking bath, counting the microcarriers in the sampled volume, and extrapolating the count to the total suspension volume. The microcarrier production rate was then calculated by dividing the estimated yield by the fabrication time.

The resulting microcarriers were collected using 100-µm cell strainers and recovered into 50-mL conical tubes by rinsing the inverted strainers with 20 mM CaCl₂ solution. Microcarrier suspensions were stored at 4 °C and used within 7 days of fabrication.

### 2.3. Layer-by-layer surface modification of microcarriers

Microcarriers were surface-modified by sequential deposition of poly-L-lysine (PLL) and tannic acid (TA). The PLL working solution contained 0.05 mg/mL PLL, 100 mM HEPES, 100 mM NaCl, and 20 mM CaCl₂, whereas the TA working solution contained 0.5 mg/mL TA, 100 mM HEPES, 100 mM NaCl, and 20 mM CaCl₂. The HEPES-containing base solution was adjusted to pH 7.0 before preparation of the coating solutions.

Microcarriers were collected using 100-µm cell strainers and recovered with 8 mL of PLL working solution. The suspension was incubated on a rotator at 30 rpm (RT-10, DAIHAN Scientific) for 15 min at room temperature, recollected using a 100-µm cell strainer, and directly resuspended in 8 mL of TA working solution without intermediate washing. After a further 15-min incubation under the same conditions, the microcarriers were collected and resuspended in 20 mM CaCl₂ solution. Single-bilayer PLL/TA-modified microcarriers were stored under refrigeration and used within 7 days.

### 2.4. Morphological characterization and microcarrier size analysis

For optical microscopy, alginate microcarriers suspended in 20 mM CaCl₂ solution were imaged using a CKX53 microscope (Olympus) equipped with a 4× objective under bright-field illumination.

For SEM analysis, bare and PLL/TA-modified alginate microcarriers were collected from 20 mM CaCl₂ storage solution using 100-µm cell strainers and washed twice with deionized water by repeated resuspension and filtration. The washed microcarriers were resuspended in 4 mL of deionized water, and a 1-mL aliquot was deposited onto carbon tape and air-dried. Samples were imaged without sputter coating using a Zeiss FESEM 1530 scanning electron microscope at an accelerating voltage of 10 kV.

Microcarrier diameters were measured from optical microscopy images using ImageJ. Microcarriers from three independent fabrication batches were analyzed, with 25 microcarriers measured per batch (75 microcarriers in total). The diameter of each microcarrier was manually measured using the straight-line tool in ImageJ after spatial calibration using the image scale bar.

### 2.5. Zeta-potential analysis during layer-by-layer assembly

Changes in microcarrier surface charge during sequential PLL and TA deposition were evaluated by zeta-potential measurements. Bare alginate microcarriers were sequentially coated with PLL and TA for three deposition cycles, yielding the following measurement points: bare alginate, +PLL(1), +TA(1), +PLL(2), +TA(2), +PLL(3), and +TA(3). After each deposition step, microcarriers were collected using 100-µm cell strainers, washed twice with deionized water, and resuspended in fresh deionized water for measurement.

Zeta potential was measured at room temperature using a Zetasizer pro (Malvern Panalytical). For each measurement, 0.8 mL of microcarrier suspension was loaded into a DTS1070 capillary cell. Three independent experiments were performed on separate days using independently prepared alginate microcarriers and PLL and TA working solutions. Data are presented as the mean of three independent experiments, with error bars indicating the minimum and maximum values.

### 2.6. Poly-L-lysine analysis by ATR-FTIR spectroscopy

ATR-FTIR spectra of bare alginate, PLL-coated alginate, and PLL/TA-coated alginate microcarriers were acquired using a Nicolet iN10 FTIR microscope (Thermo Fisher Scientific, USA) equipped with a germanium attenuated total reflectance (ATR) tip. Microcarriers were air-dried, removed from the drying substrate, and directly analyzed without further grinding. Spectra were acquired over 400–4000 cm⁻¹ using 128 scans per spectrum. For spectral comparison, each spectrum was individually normalized to a 0–1 range using min–max normalization.

### 2.7. Tannic acid analysis by Prussian Blue assay and UV-Vis spectroscopy

Phenolic content associated with bare alginate, PLL-coated alginate, and PLL/TA-coated alginate microcarriers was assessed using the Prussian Blue assay (Joshua et al., 2016). The Prussian Blue reagent was freshly prepared by mixing equal volumes of 2% (w/v) FeCl₃ and 2% (w/v) potassium ferricyanide solutions. For each condition, 2 mL of microcarrier suspension was collected using a 100-µm cell strainer and recovered in 2 mL of deionized water. After incubation at room temperature for at least 10 min, the suspension was centrifuged at 4,000 × g for 1 min and filtered again. The microcarriers were subsequently washed four times with 2 mL of deionized water. After the final wash, the filtrate was collected as the indirect sample, and microcarriers retained on the strainer were recovered in 2 mL of deionized water as the direct sample.

For the assay, 200 µL of each sample was mixed with 10 µL of Prussian Blue reagent and transferred to a transparent 96-well plate. Deionized water containing the same volume of reagent was used as the blank. Samples were incubated at room temperature for 30 min, and absorbance was measured at 750 nm using a Spark® 10M microplate reader (Tecan, Männedorf, Switzerland).

### 2.8. Cell culture

MDA-MB-231 cells were maintained in DMEM supplemented with 10% FBS. Hanwoo bovine and Handon porcine muscle satellite cells were maintained in DMEM supplemented with 10% FBS, 1% P/S, and 2% growth factor (SeaWith, South Korea). The supplier-reported characteristics and authentication methods of the livestock-derived cells are summarized in Supplementary Table S1. All cells were cultured at 37 °C in a humidified atmosphere containing 5% CO₂.

### 2.9. Cell seeding and culture on microcarriers

For the alginate model system, MDA-MB-231 cells were seeded onto bare alginate, PLL-coated alginate, TA-treated alginate, and PLL/TA-modified alginate microcarriers. For cross-polysaccharide evaluation, Hanwoo bovine and porcine muscle cells were cultured on bare and PLL/TA-modified pectin and carrageenan microcarriers.

For each condition, microcarriers were collected from 4 mL of microcarrier suspension using a 100-μm cell strainer and transferred to a 6-well plate by rinsing the inverted strainer with 4 mL of the corresponding growth medium. Cells harvested from a confluent T75 flask were resuspended in 4 mL of growth medium, and 0.3 mL of the cell suspension, corresponding to approximately 6.3 × 10⁵ cells, was added to each well to give a final culture volume of 4.3 mL. The suspensions were gently mixed by pipetting to promote cell–microcarrier contact and subsequently maintained under static culture conditions.

For the MDA-MB-231 alginate model system, cultures were maintained for 6 days without medium replacement. For Hanwoo bovine and porcine muscle cell experiments, cultures were maintained for up to 6 days without medium replacement, with analyses performed on Days 1, 3, and 6.

### 2.10. Live-cell staining and confocal imaging

Cell association with microcarriers was visualized using Calcein-AM staining, while propidium iodide was used to identify membrane-compromised cells. A 10 mM Calcein-AM stock solution was prepared in DMSO and diluted to a final concentration of 1.25 µM. Samples were stained for 30 min at 37 °C. Propidium iodide (PI) was added at a final concentration of 1.875 µM. Fluorescence images were acquired using a STELLARIS 5 confocal microscope (Leica Microsystems) equipped with a 10x and 20x objectives.

For the MDA-MB-231 alginate model system, separate samples were imaged from Day 1 to Day 6. Hanwoo and porcine muscle-cell cultures were imaged at Day 1, 3, and 6.

The same Calcein-AM/PI staining and confocal imaging procedure was applied to Hanwoo bovine and porcine muscle cell cultures.

### 2.11. Quantification of cell coverage and initial attachment

Calcein-positive cell coverage in the MDA-MB-231 alginate model system was quantified from fluorescence images using ImageJ. The projected microcarrier area and Calcein-positive area were measured using intensity thresholding followed by area measurement. Cell coverage was calculated as the Calcein-positive area divided by the projected microcarrier area and expressed as a percentage (Fig. S5). Measurements were performed in three independent experiments for each condition and time point.

Initial cell attachment in the alginate model system was evaluated from Day 1 images. Microcarriers containing at least one attached Calcein-positive cell were classified as cell-bearing, and the percentage of cell-bearing microcarriers was calculated relative to the total number of microcarriers analyzed. The number of attached cells per microcarrier was determined by manual counting of individual Calcein-positive cells and averaged across all analyzed microcarriers, including those without attached cells. Analysis was performed across three independent experimental batches, with 20 microcarriers analyzed per batch (60 microcarriers in total) for each condition.

For Hanwoo bovine and porcine muscle cells cultured on pectin and carrageenan microcarriers, cell coverage was quantified using the same image-analysis procedure. However, microcarriers with at least 2% Calcein-positive cell coverage were classified as cell-bearing for the Day 1 attachment analysis. Ten microcarriers were analyzed per condition from a single experiment, and cell-covered area was quantified individually for each microcarrier. For temporal cell-coverage analysis, three microcarriers were analyzed per condition at each time point (Days 1, 3, and 6) from the same experimental batch.

### 2.12. Protein adsorption quantification by NanoOrange assay

Protein adsorption onto microcarriers was evaluated using the NanoOrange Protein Quantitation Kit (Thermo Fisher Scientific) according to the manufacturer’s instructions. For each condition, microcarriers were collected from 4 mL of microcarrier suspension using a 100-μm cell strainer and transferred to a 6-well plate by rinsing the inverted strainer with 4 mL of growth medium (GM). The microcarrier suspensions were incubated at 37 °C for 10 min with shaking to allow protein adsorption from the growth medium. Following incubation, the suspensions were centrifuged at 6,000 × g for 1 min to sediment the microcarriers, and the supernatants were collected for protein quantification. Protein quantification was performed using the NanoOrange assay according to the manufacturer’s instructions. Fluorescence intensity was measured in 96-well flat-bottom black plates (Greiner Bio-One) using a SPARK CYTO multimode microplate reader (Tecan, Switzerland) at excitation and emission wavelengths of 485 and 535 nm, respectively. Protein concentrations were determined using a standard curve generated with bovine serum albumin (BSA). The amount of protein adsorbed onto the microcarriers was estimated from the difference in protein concentration between the original growth medium and the supernatant after incubation with microcarriers. Two independent experiments were performed for each condition (n = 2).

Statistical significance between bare and PLL/TA-modified alginate microcarriers was assessed using Student’s t-test.

### 2.13. Fluorescence analysis of surface-associated fibronectin

Surface association of fibronectin with bare alginate and PLL/TA-modified alginate microcarriers was evaluated using fluorescently labeled fibronectin. HiLyte™ 488-labeled fibronectin (FNR02; Cytoskeleton, Inc., Denver, CO, USA) was prepared in DMEM at a concentration of 50 μg/mL.

For each condition, 200 μL of microcarrier suspension was collected using a 100-μm cell strainer. The retained microcarriers were recovered by rinsing the inverted strainer with 200 μL of fibronectin-containing DMEM. The resulting microcarrier suspensions were incubated on a shaker at 37 °C for 10 min.

Following incubation, microcarrier-associated fluorescence was visualized by confocal fluorescence microscopy using the Alexa Fluor 488 detection channel. All samples were imaged using identical confocal acquisition settings.

Fluorescence images were analyzed using ImageJ software (National Institutes of Health, USA). Z-stack images were converted to maximum-intensity projections prior to analysis. The entire microcarrier was defined as the region of interest (ROI), and the mean fluorescence intensity with each ROI was measured. Background-corrected fluorescence intensity was calculated by subtracting the mean background fluorescence intensity from the mean fluorescence intensity of each microcarrier ROI. Five individual microcarriers were analyzed per condition (n = 5 microcarriers).

Statistical significance in background-corrected fluorescence intensity between bare and PLL/TA-modified alginate microcarriers was assessed using Student’s t-test.

## 3. Results and Discussion

To implement the decoupled surface-engineering strategy, we established a three-step scheme comprising microcarrier fabrication, surface modification, and protein recruitment (Fig. 1). Using alginate as a representative polysaccharide, an alginate solution was first sprayed into a calcium-containing solution to generate microsized droplets, which rapidly gelled through Ca²⁺-mediated crosslinking of the alginate chains (Step 1). This spray-based process enabled facile production of microsized microcarriers before introducing cell-interactive surface functionality. The preformed microcarriers were sequentially treated with PLL and TA to construct an interface combining electrostatic assembly on anionic polysaccharide surfaces with the protein-interacting properties of TA (Step 2). This post-fabrication modification allowed the surface properties to be engineered independently of the underlying microcarrier composition. Upon exposure to the culture environment, the PLL/TA-modified interface recruited medium-derived proteins onto the microcarrier surface (Step 3), allowing cell-interactive functionality to emerge during culture without predefined protein presentation. Together, this three-step strategy decouples bulk microcarrier fabrication from cell-interactive surface functionality, providing a modular framework for edible microcarrier design.

**Figure 1.**
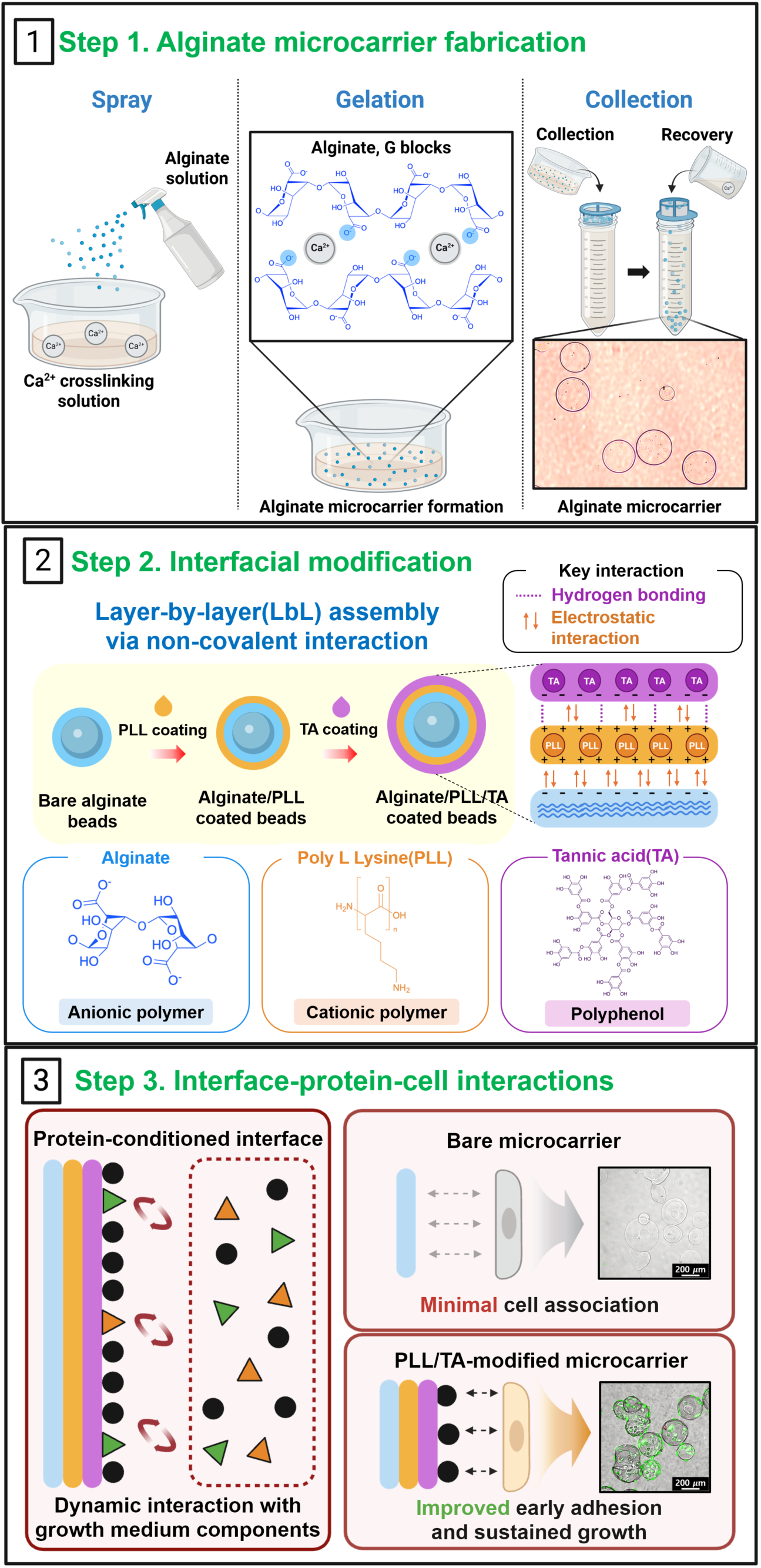
Decoupled surface-engineering strategy for introducing cell-interactive functionality to edible polysaccharide microcarriers. (1) Alginate microcarriers are generated by spraying an alginate solution into a Ca²⁺-containing bath, followed by rapid Ca²⁺-mediated gelation. (2) The preformed microcarriers are sequentially modified with poly-L-lysine (PLL) and tannic acid (TA) to construct a PLL/TA interface independently of the underlying microcarrier composition. (3) The PLL/TA-modified interface recruits proteins from the culture environment, allowing cell-interactive functionality to emerge during culture without predefined protein presentation.

We first established spray-based Ca²⁺ gelation as a facile method for producing large populations of alginate microcarriers with consistent morphology and size. A 2.5% (w/v) alginate solution was sprayed into a 20 mM CaCl₂ solution at a flow rate of 22.42 mL min⁻¹, generating microsized droplets that rapidly gelled through Ca²⁺-mediated crosslinking of the alginate chains. Under these conditions, approximately 95,000 microcarriers were produced within 0.89 min, corresponding to a production rate of approximately 1.07×10^5^ microcarriers min⁻¹. Optical microscopy revealed large populations of discrete, similarly sized microcarriers without obvious aggregation (Fig. 2A). Across three independently prepared batches, median diameters ranged from 149 to 187 μm, with interquartile ranges of 130–190, 137–214, and 136–222 μm, respectively (Fig. 2B). The substantial overlap among these distributions indicates reproducible generation of comparable microcarrier populations across independent fabrication runs.

**Figure 2.**
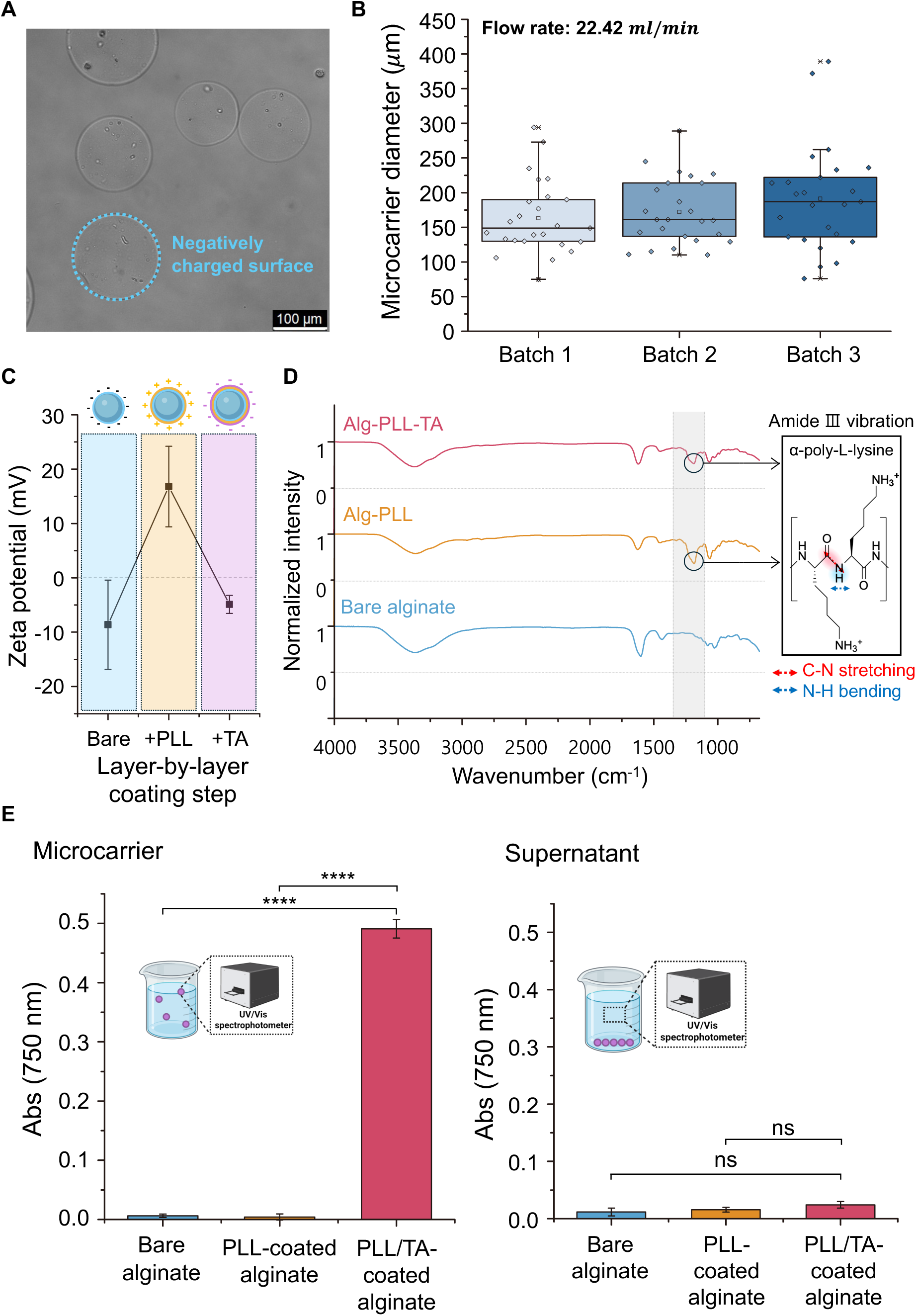
Fabrication and post-fabrication surface modification of alginate microcarriers with PLL and TA. (A) Representative optical micrograph of spray-fabricated alginate microcarriers. Scale bar, 100 μm. (B) Diameter distributions across three independently prepared batches (n = 25 microcarriers per batch). Center lines indicate medians; boxes indicate interquartile ranges. (C) Zeta potential during sequential PLL and TA treatment. Values are mean with minimum and maximum values (n = 3 independent experiments). (D) ATR-FTIR spectra of bare alginate, PLL-treated, and PLL/TA-treated microcarriers, individually min–max normalized to 0–1. The highlighted 1220–1330 cm⁻¹ region denotes the amide III region. (E) Prussian blue analysis of microcarriers and corresponding supernatants. Absorbance was measured at 750 nm. Values are mean ± SD (n = 3). Statistical comparisons were performed using the Tukey–Kramer multiple-comparisons test. ns, not significant; ****p < 0.0001.

We next examined whether a PLL/TA interface could be sequentially constructed on the preformed alginate microcarriers. SEM images showed qualitative differences in surface morphology between bare and PLL/TA-modified microcarriers, with the modified microcarriers appearing more heterogeneous and exhibiting particulate or clustered surface features (Fig. S1). Zeta-potential measurements demonstrated alternating changes in surface charge following sequential PLL and TA treatment, with the initially negative alginate surface shifting from −8.6 mV to +16.8 mV after PLL treatment and returning to −4.9 mV after subsequent TA treatment (Fig. 2C). The alternating surface charge pattern persisted through subsequent coating cycles (Fig. S4). Because these electrokinetic measurements do not directly identify coating composition, chemical characterization was used to further assess the modified interface. Attenuated total reflectance Fourier-transform infrared spectroscopy (ATR-FTIR) revealed an additional feature in the 1220–1330 cm⁻¹ amide III region after PLL treatment, consistent with PLL association (Mirtič & Grdadolnik, 2013), whereas a distinct TA-specific band could not be clearly resolved (Fig. 2D). Complementary Prussian blue analysis yielded a markedly greater phenolic response in PLL/TA-treated microcarriers than in bare alginate and PLL-treated controls, while the corresponding supernatants exhibited low absorbance with no significant differences among groups (Fig. 2E). These results support the retention of TA-associated phenolic material on the microcarriers after washing, with no significant accumulation of detectable phenolic material in the supernatant under the tested conditions. Together, these morphological, electrokinetic, spectroscopic, and phenolic measurements support the sequential construction of a PLL/TA interface on preformed alginate microcarriers, demonstrating that surface engineering can be implemented independently of the initial microcarrier fabrication process.

Using MDA-MB-231 cells and alginate microcarriers as a model system, PLL/TA modification markedly enhanced initial cell attachment. At Day 1, cells were readily observed on PLL/TA-modified microcarriers, whereas bare alginate and TA-treated microcarriers exhibited little detectable cell attachment, and PLL-coated microcarriers supported only limited attachment (Fig. 3A). Quantitative analysis revealed that 83% of PLL/TA-modified microcarriers supported at least one attached cell, compared with 36% of PLL-coated microcarriers and no detectable attachment on either bare alginate or TA-treated microcarriers (Fig. 3B). PLL/TA-modified microcarriers also supported an average of 2.95 attached cells per microcarrier, compared with 0.85 on PLL-coated alginate. These findings indicate that the combined PLL/TA interface provided a substantially greater adhesion-promoting effect than either component alone.

**Figure 3.**
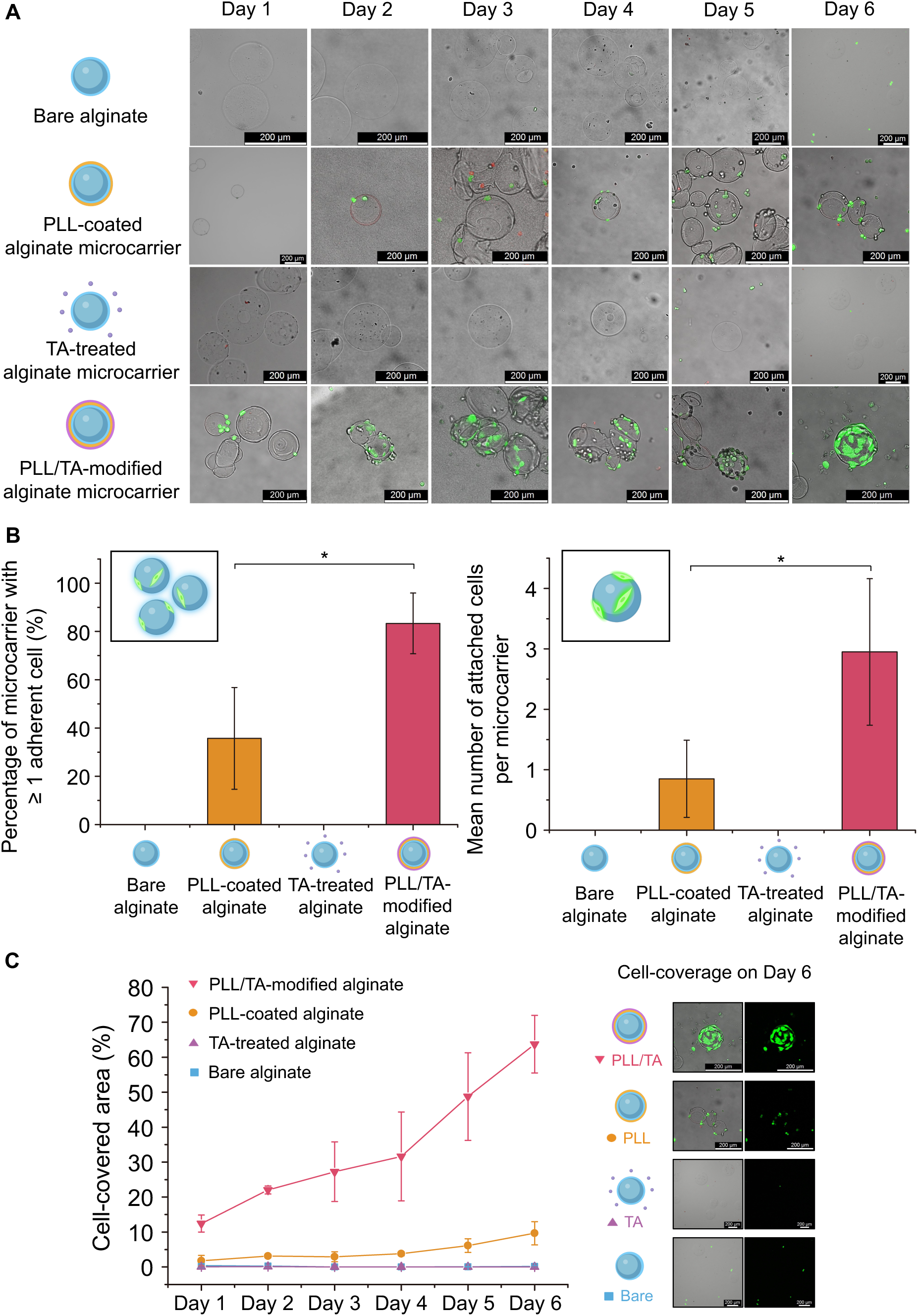
PLL/TA interface engineering enhances initial cell attachment and subsequent cell expansion on alginate microcarriers. (A) Representative confocal fluorescence images of Calcein-AM-stained MDA-MB-231 cells cultured on bare alginate, PLL-coated, TA-treated, and PLL/TA-modified microcarriers from Day 1 to Day 6. (B) Initial cell attachment at Day 1, expressed as the percentage of microcarriers bearing at least one attached cell and the mean number of attached cells per microcarrier. Values represent mean ± SD from three independent experiments with 20 microcarriers analyzed per batch (60 microcarriers in total). (C) Cell-covered area from Day 1 to Day 6, calculated as the Calcein-positive area divided by the projected microcarrier area after intensity thresholding in ImageJ. Values represent mean ± SD from three independent experiments for each condition and time point.

The difference between PLL/TA-modified and control microcarriers became more pronounced during culture. Fluorescence imaging revealed progressively increasing cell coverage on PLL/TA-modified microcarriers, while the other conditions remained sparsely populated (Fig. 3A). By Day 6, cell-covered area reached 63.74% on PLL/TA-modified microcarriers compared with 9.64% on PLL-coated microcarriers, whereas cell coverage remained negligible on bare alginate and TA-treated controls (Fig. 3C). Together, these results demonstrate that PLL/TA interface engineering transformed poorly adhesive alginate microcarriers into cell-supportive surfaces that promoted both initial cell attachment and subsequent cell expansion.

Following the enhanced cell attachment and coverage observed on PLL/TA-modified alginate microcarriers, we investigated the interaction between the modified interface and proteins in the culture environment. Tannic acid is known to interact with proteins through multivalent noncovalent interactions (Jafari et al., 2022), while protein adsorption from serum-containing media onto polysaccharide-based scaffolds has also been reported (Nga et al., 2025). Based on these properties, we proposed that PLL/TA modification could promote protein association at the microcarrier surface, potentially including proteins relevant to cell adhesion, as illustrated schematically in Fig. 4A.

**Figure 4.**
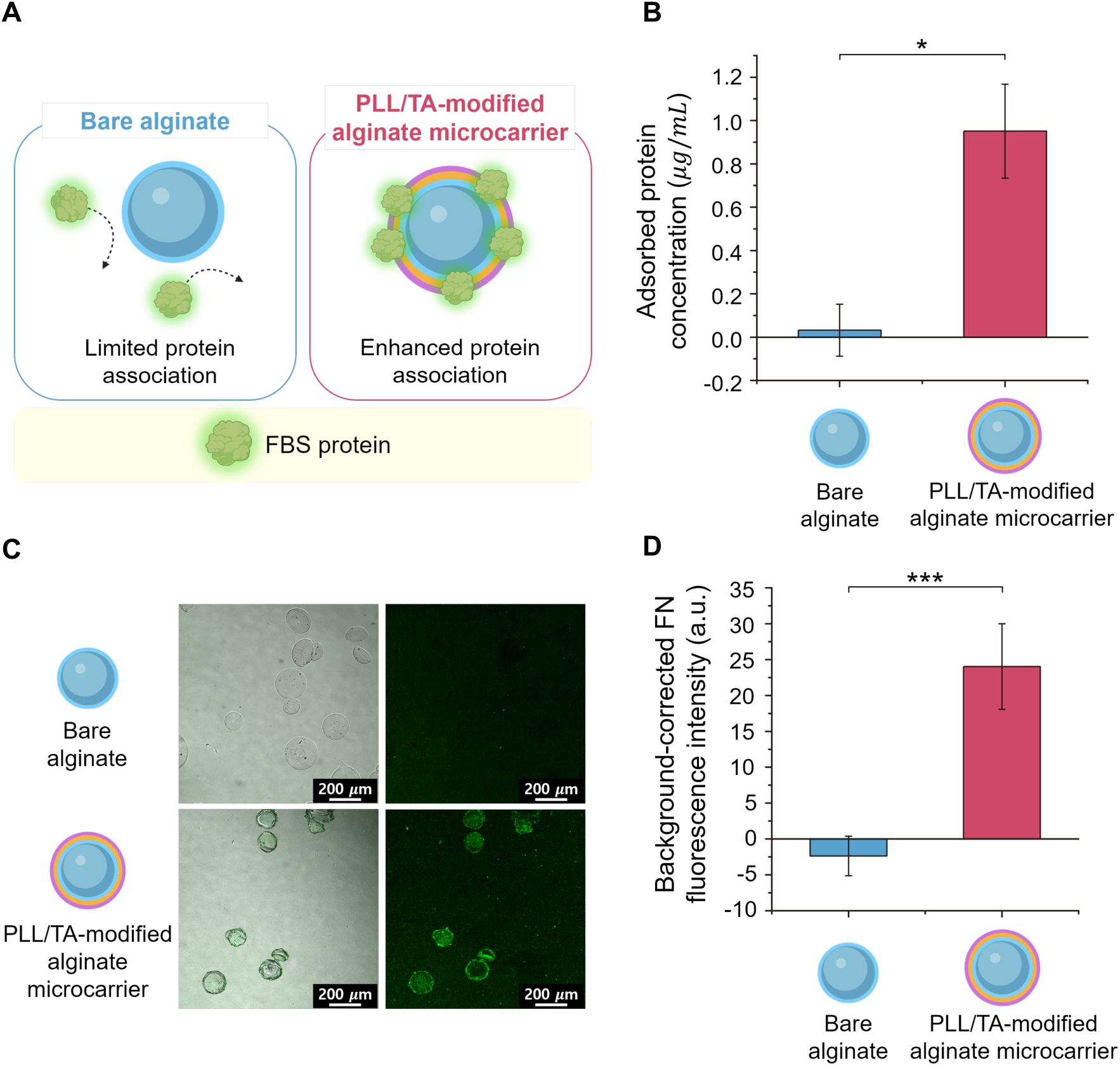
PLL/TA modification promotes the recruitment of culture-medium proteins onto alginate microcarriers. (A) Schematic illustration of protein association with bare and PLL/TA-modified alginate microcarriers. (B) Estimated protein association with microcarriers, calculated from the decrease in soluble protein concentration following incubation with growth medium. Values are mean ± SD (n = 2 independent experiments); *p < 0.05. (C) Representative confocal fluorescence images of fibronectin (FN) associated with bare and PLL/TA-modified alginate microcarriers. Scale bars, 200 μm. (D) Background-corrected FN fluorescence intensity. Values are mean ± SD (n = 5 technical replicates); ***p < 0.001. Statistical comparisons in (B) and (D) were performed using Student’s t-test.

To evaluate this possibility, protein association was estimated using a NanoOrange assay by measuring protein depletion from growth medium following incubation with microcarriers. Protein concentrations were determined using a BSA calibration curve (Fig. S2). Estimated protein association was 0.95 ± 0.22 μg/mL for PLL/TA-modified microcarriers, compared with 0.03 ± 0.12 μg/mL for bare alginate controls (Fig. 4B). To assess whether this protein-associating capability extended to adhesion-related proteins, we next examined fibronectin (FN), a representative extracellular-matrix protein involved in cell adhesion (Winkler et al., 2020). Confocal imaging revealed greater FN-associated fluorescence on PLL/TA-modified microcarriers than on bare alginate controls (Fig. 4C). Background-corrected fluorescence intensity reached 23.74 ± 6.29 a.u. for PLL/TA-modified microcarriers, compared with −2.37 ± 2.76 a.u. for bare controls (Fig. 4D).

The enhanced association of both culture-medium proteins and FN demonstrates the protein-recruiting capability of the PLL/TA-modified interface. This property offers a potential route for introducing cell-interactive functionality through interactions with the culture environment, rather than relying on the predefined presentation of specific adhesion-active proteins.

To evaluate the transferability of PLL/TA interface engineering beyond alginate, we selected pectin and carrageenan as additional food-grade polysaccharides that share two relevant characteristics with alginate, anionic functional groups and the ability to undergo Ca²⁺-mediated gelation (Capel et al., 2006; Kalsi et al., 2025). We applied the same surface modification to microcarriers fabricated from these polysaccharides and investigated their interactions with Hanwoo bovine and Handon porcine muscle satellite cells (Fig. 5A).

**Figure 5.**
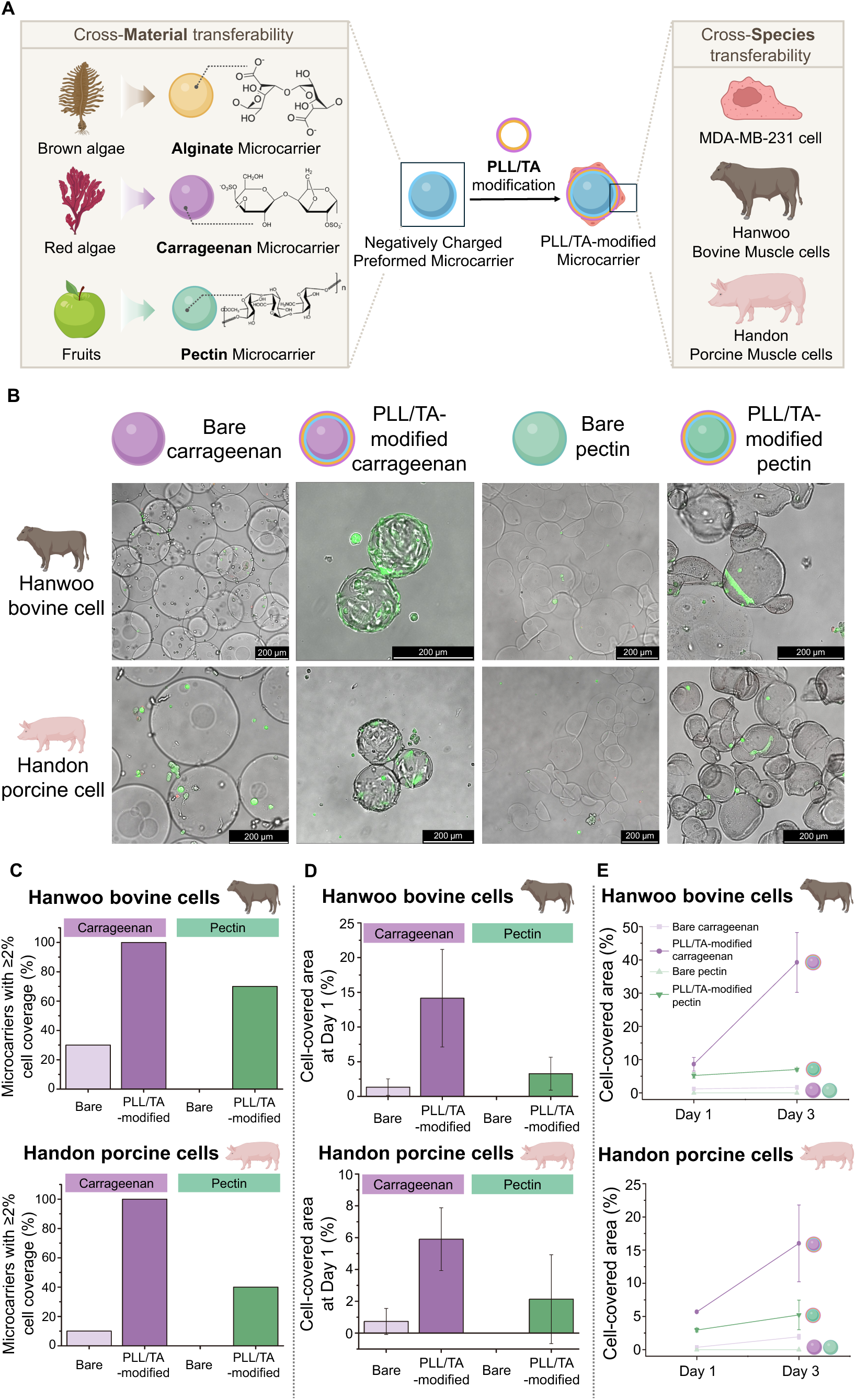
A decoupled PLL/TA interface-engineering strategy is transferable across edible polysaccharide microcarriers and livestock-derived cell sources. (A) Schematic illustrating the application of PLL/TA interface engineering across different edible polysaccharides (alginate, carrageenan, and pectin) and cell sources (MDA-MB-231, Hanwoo bovine, and Handon porcine muscle cells). (B) Representative confocal fluorescence images showing Hanwoo bovine and Handon porcine muscle cells attached to PLL/TA-modified carrageenan and pectin microcarriers at Day 1. (C) Initial cell attachment at Day 1, expressed as the percentage of microcarriers with at least 2% cell-covered area. (D) Cell-covered area at Day 1, calculated as the Calcein-positive area divided by the projected microcarrier area. (E) Changes in cell-covered area during early culture (Days 1-3) for Hanwoo bovine and Handon porcine muscle cells cultured on bare and PLL/TA-modified carrageenan and pectin microcarriers. In panels C and D, n = 10 microcarriers per condition from a single experiment. In panel E, values are mean ± SD from n = 3 microcarriers per condition and time point from a single experiment.

Representative confocal images revealed cell–microcarrier association across the different PLL/TA-modified polysaccharide–livestock cell combinations, showing that both bovine and porcine muscle cells could interact with the modified carrageenan and pectin surfaces (Fig. 5B). Quantitative analysis demonstrated that PLL/TA modification enhanced initial cell attachment across all four tested combinations. The proportion of microcarriers exhibiting at least 2% cell-covered area increased from 30% to 100% for Hanwoo bovine cells on carrageenan and from 0% to 70% on pectin, while the corresponding values for Handon porcine cells increased from 10% to 100% and from 0% to 40%, respectively (Fig. 5C). Consistent with these results, a separate Day 1 experiment revealed greater cell-covered area on PLL/TA-modified microcarriers than on the corresponding bare controls across both polysaccharides and cell sources (Fig. 5D). Although the magnitude of the response differed among material–cell combinations, the same PLL/TA modification consistently promoted early cell attachment across all tested conditions.

In an independent time-course experiment, PLL/TA-modified microcarriers exhibited greater cell-covered area than their corresponding bare controls during the first three days of culture across all four polysaccharide–cell combinations (Fig. 5E). On modified carrageenan, for example, cell-covered area increased from 8.61% to 39.2% for Hanwoo bovine cells and from 5.70% to 16.0% for Handon porcine cells between Days 1 and 3. However, subsequent changes in cell coverage varied among material–cell combinations, with some showing declines after Day 3 (Fig. S3). These findings indicate that PLL/TA modification enhances early cell coverage, although sustained cell expansion remains dependent on the specific polysaccharide–cell combination, consistent with previous observations that cellular responses to engineered interfaces can be influenced by the underlying substrate (Keselowsky et al., 2003; Kim et al., 2012).

Taken together with the alginate model system, these findings demonstrate that the early adhesion-promoting effect of PLL/TA modification extends across different edible polysaccharides and cell sources, while sustained cell coverage remains dependent on the specific material–cell combination. More broadly, this decoupled strategy enables cell-interactive surface functionality to be introduced onto preformed microcarriers without changing their bulk formulation, expanding the range of edible polysaccharides that can be considered for cultivated meat applications.

## Conclusion

This study establishes a decoupled surface-engineering strategy for introducing cell-interactive functionality onto preformed polysaccharide microcarriers without predefined adhesion-active protein presentation. On alginate microcarriers, PLL/TA modification increased the proportion of microcarriers supporting initial cell attachment from 0% on bare controls to 83%. By Day 6, cell-covered area reached 63.74%, compared with 9.64% on PLL-coated controls. The modified interface also promoted the recruitment of culture-medium proteins and the association of fibronectin, consistent with an adaptive mechanism in which cell-supportive functionality can emerge through protein recruitment. Importantly, the adhesion-promoting effect extended beyond the alginate model system to pectin and carrageenan microcarriers cultured with Hanwoo bovine and Handon porcine muscle cells.

Beyond the observed improvements in cell attachment, this work introduces a more flexible approach to edible microcarrier design. By separating surface functionalization from bulk fabrication, different polysaccharides can be considered for their food-related properties without incorporating predefined adhesion-supportive proteins into each formulation. The variation in longer-term cellular responses nevertheless underscores that this design flexibility does not eliminate the influence of the underlying material and cell source.

Further studies should define the composition and functional roles of the recruited protein layer, optimize the interface for specific material–cell combinations, and evaluate long-term cell expansion under scalable dynamic culture conditions. Molecular recognition and site-selective conjugation strategies could further enable selective protein recruitment and controlled functionalization of recruited proteins (Jo et al., 2025; Kim et al., 2023), while advanced biopolymer patterning techniques could extend this approach beyond spherical microcarriers to more complex three-dimensional architectures (Yang et al., 2023). Validation of food-related properties, including texture, nutritional contribution, and processing compatibility, will also be essential for translating these microcarriers into cultivated meat production. Ultimately, this strategy expands the design space for edible microcarriers by allowing food-grade polysaccharides to be selected for their intended roles in the final product while cell-interactive functionality is engineered separately at the interface.

## Acknowledgement

This work was supported by the National Research Foundation of Korea (NRF) grant funded by the Ministry of Science and ICT (RS-2026-25605637, RS-2025-16070008, and RS-2025-25401663), the Korea Health Industry Development Institute grant funded by the Ministry of Health and Welfare (RS-2024-00507436), the Commercialization Promotion Agency for R&D Outcomes grant funded by the Ministry of Science and ICT (RS-2026-25608867), and the Korea Basic Science Institute (National Research Facilities and Equipment Center) grant funded by the Ministry of Education (RS-2026-25539607). This work was partially supported by the Program funded by the Gyeongsangbuk-do and Pohang-si (Synthetic Biology grant).

