## Supplementary Information for "Decoupled Surface Engineering Enables Cell-Interactive Edible Microcarriers across Food-Grade Polysaccharides"

### Supplementary Figures

#### Supplementary Figure S1

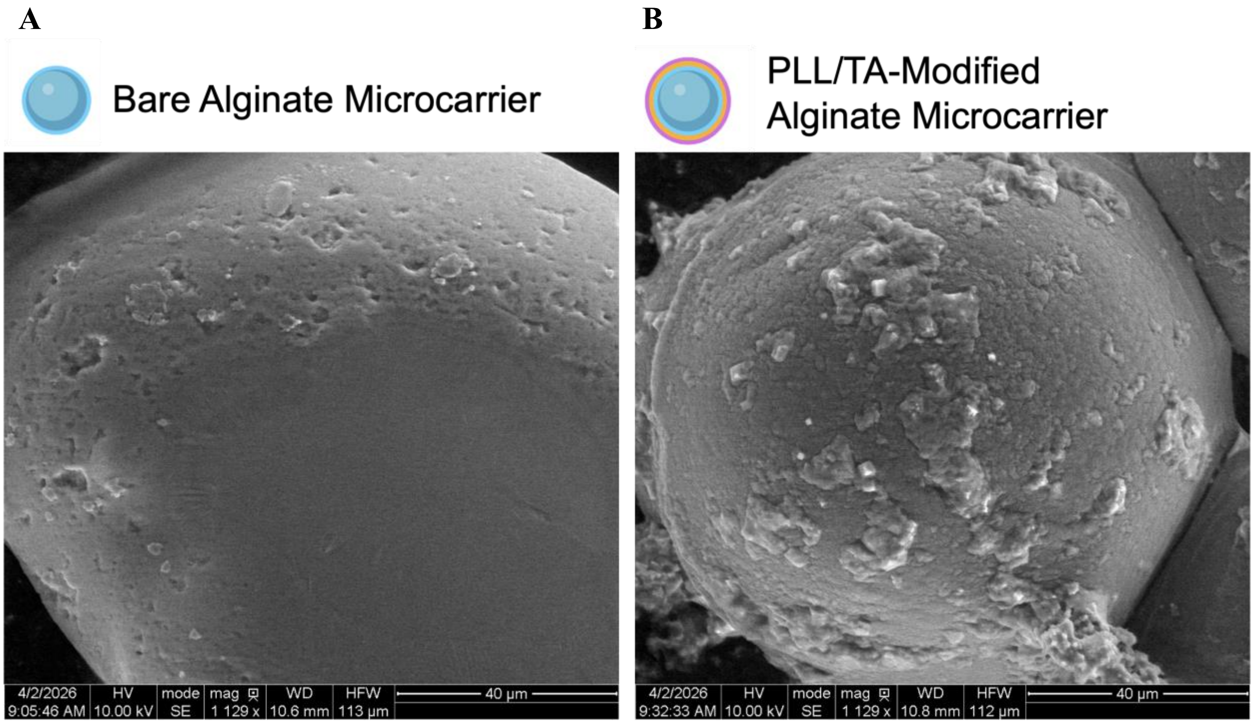


**Supplementary Figure S1. Representative scanning electron microscopy (SEM) images of bare alginate and PLL/TA-treated microcarriers.**

(A) Bare alginate microcarrier and (B) PLL/TA modified alginate microcarrier. The modified exhibits a more heterogeneous surface appearance with particulate and clustered features.

**Supplementary Figure S2**

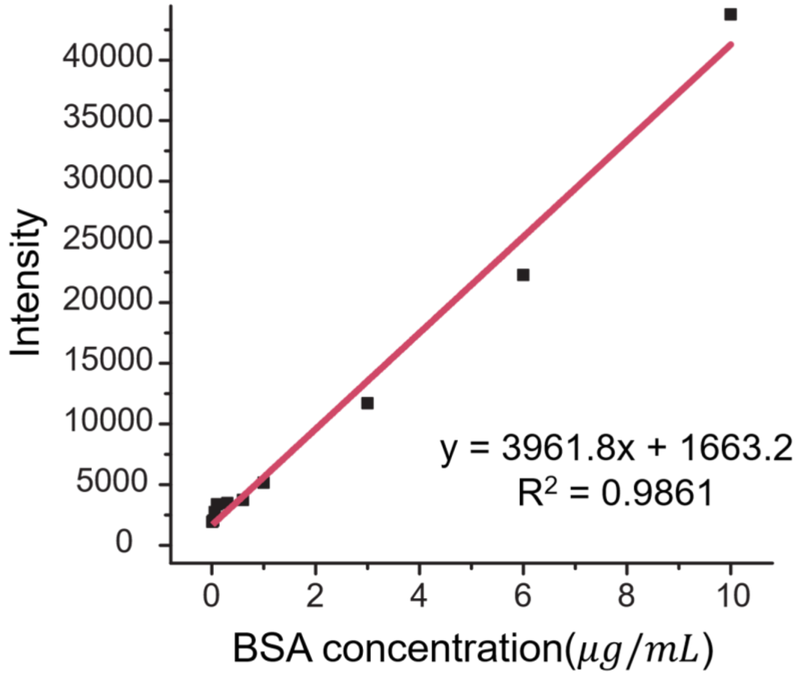


**Supplementary Figure S2. BSA calibration curve for NanoOrange protein quantification.** Fluorescence intensity measured using the NanoOrange assay at different BSA concentrations. The solid line represents a linear regression fit used to convert fluorescence intensity into protein concentration for estimating protein depletion from growth medium following microcarrier incubation.

**Supplementary Figure S3**

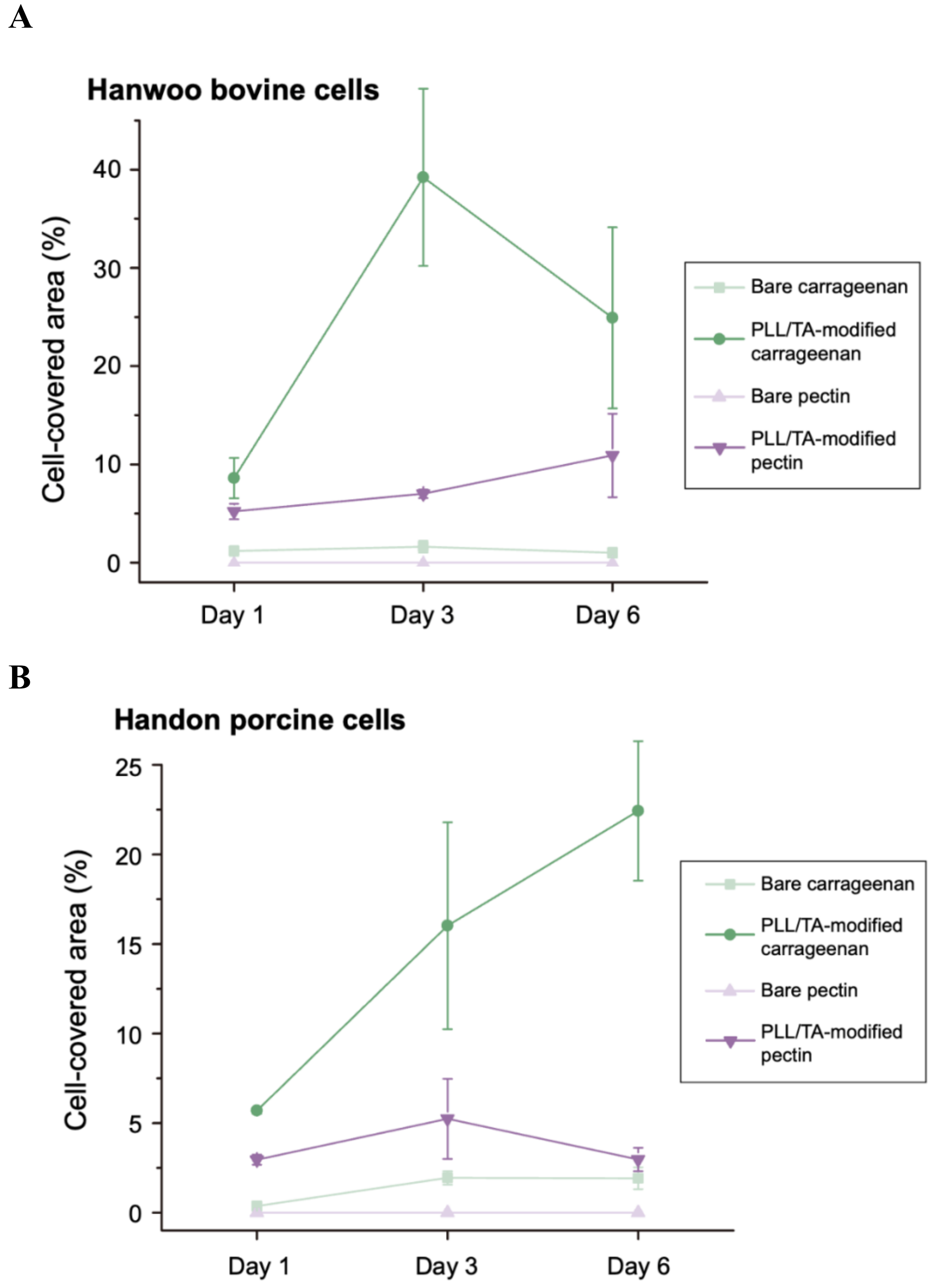


**Supplementary Figure S3. Time-dependent changes in cell coverage on bare and PLL/TA-modified polysaccharide microcarriers during six-day culture.** Cell-covered area of (A) Hanwoo bovine and (B) Handon porcine muscle cells cultured on bare and PLL/TA-modified carrageenan and pectin microcarriers at Days 1, 3, and 6. Cell coverage was calculated as the Calcein-positive area divided by the projected microcarrier area. Values are mean ± SD (n = 3 microcarriers per condition and time point from a single experiment).

**Supplementary Figure S4**

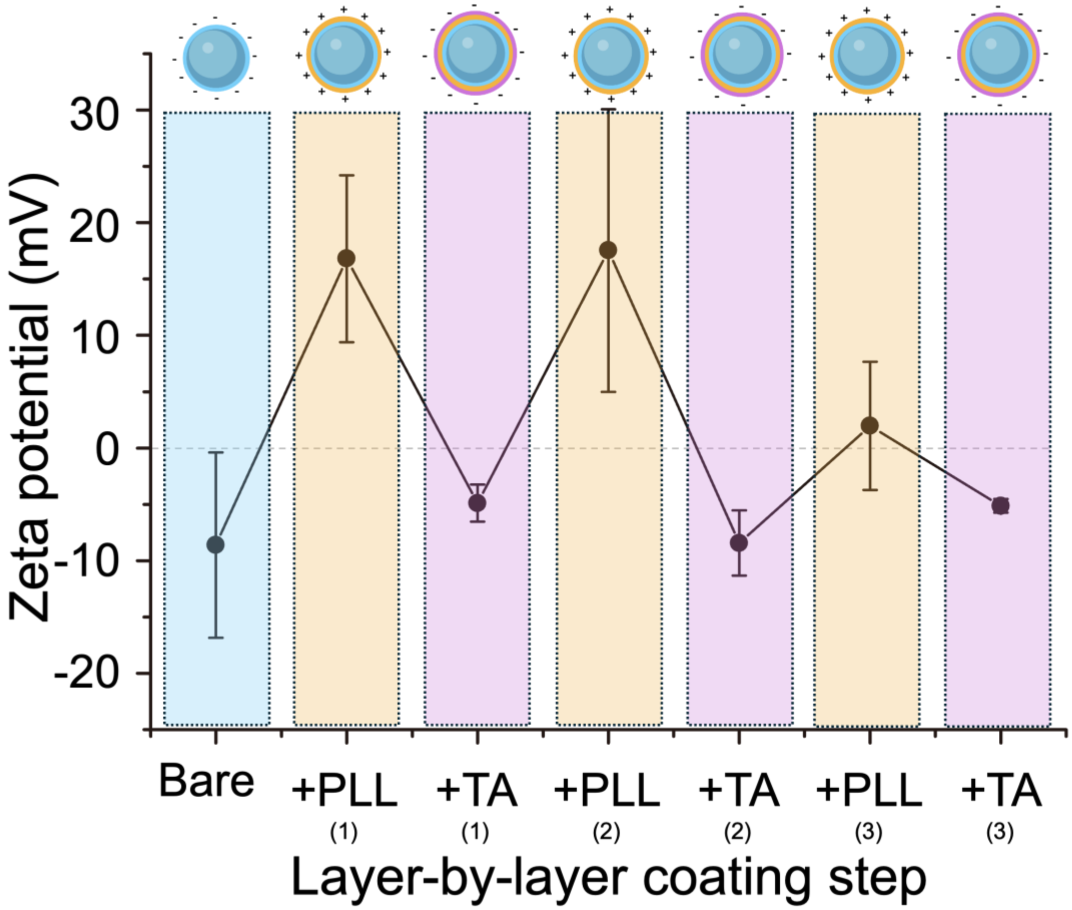


**Supplementary Figure S4. Zeta potential changes during sequential layer-by-layer coating of alginate microcarriers.** Zeta potential measurements of bare alginate microcarriers and microcarriers following successive deposition of poly-L-lysine (PLL) and tannic acid (TA) over three coating cycles. Alternating positive and negative zeta potentials were observed after PLL and TA deposition, respectively, supporting sequential surface modification.

**Supplementary Figure S5**

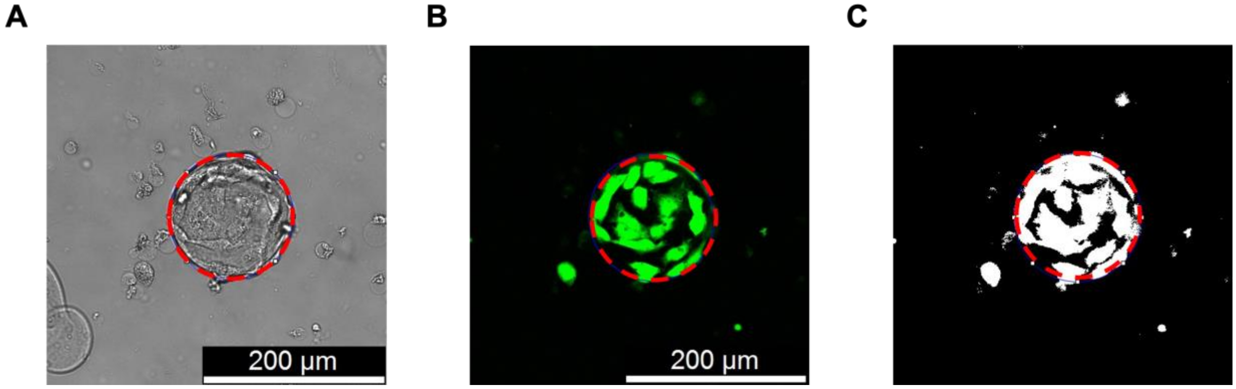


**Supplementary Figure S5. Quantification of cell-covered area on microcarriers.** Representative images of (A) bright-field microscopy, (B) confocal fluorescence microscopy of Calcein-AM-stained cells, and (C) the corresponding thresholded fluorescence image with the projected microcarrier area defined as the region of interest (ROI). A consistent intensity threshold was applied to the Calcein-AM channel using ImageJ to identify Calcein-AM-positive regions. The cell-covered area (%) was quantified using the Area Fraction (% Area) measurement within the defined ROI.

**Supplementary Table S1. Characteristics and authentication information of bovine and porcine muscle satellite cells used in this study.**

| | **Bovine cells** | **Porcine cells** |
|---|---|---|
| **Species** | *Bos taurus coreanae* | *Sus scrofa* |
| **Reported breed** | Hanwoo | Handon |
| **Cell type** | Muscle satellite cell | Muscle satellite cell |
| **Tissue origin** | Skeletal muscle | Skeletal muscle |
| **Growth properties** | Adherent | Adherent |
| **Supplier** | SeaWith Inc. | SeaWith Inc. |
| **Product/Lot ID** | Cell code: 790512 | Cell code: D24171 |
| **Authentication method** | MC1R analysis and STR profiling | Species-specific DNA testing and STR profiling |
